# Human pluripotent stem cell-derived macrophages modify development of human kidney organoids

**DOI:** 10.1101/2025.11.07.687192

**Authors:** Filipa M. Lopes, Ioannis Bantounas, Alexandra Sarov, Adrian S. Woolf, Susan J. Kimber

**Affiliations:** Division of Cell Matrix Biology and Regenerative Medicine, Faculty of Biology, Medicine and Health, University of Manchester, and the Manchester Academic Health Science Centre, Manchester, UK

**Keywords:** fetus, glomerulus, kidney, macrophage, organoid, stem cell, vessel

## Abstract

**Introduction:** The human fetal kidney contains macrophages, innate immune cells postulated to enhance its development. Macrophages have also been implicated in the pathobiology of human kidney malformations and Wilms tumour. Human pluripotent stem cell (hPSC)-derived kidney organoids contain nephrons differentiated from intermediate mesoderm-like precursors. Endothelia are present between organoid tubules but fail to efficiently populate glomeruli. These organoids lack macrophages, as expected because *in vivo* kidney macrophages invade from yolk sac and liver.

**Methods:** We hypothesised that combining hPSC-derived macrophages with hPSC-derived kidney precursors modifies nephrogenesis. Macrophages harvested at early or later maturation stages were added to kidney precursors in numbers of 1%, 5% or 20% compared with constant numbers of nephrogenic cells.

**Results:** No macrophages, as assessed by CD68 immunostaining, were detected in organoids without added macrophages. In contrast, composite organoids contained macrophages located between tubules, mimicking native human fetal kidneys. Quantification of tissue macrophages at the end of 18-day organoid culture positively correlated with the numbers of macrophages added. Assessed by CD31/PECAM-1 and CD68 co-immunostaining, some macrophages were near vessels but added macrophages neither increased the proportion of vessels nor endothelial invasion of glomeruli. Early-stage macrophages significantly increased the percentage area occupied by glomeruli, as assessed by synaptopodin immunostaining. The highest macrophage numbers inhibited overall growth, assessed by organoid area, additionally generating dysmorphic tissue when later stage macrophages were added.

**Conclusion:** Depending on their maturation stage and quantity, macrophages have beneficial or harmful effects on organoids. This supports the proposition that macrophages play roles in normal and abnormal development of human kidneys.

## Introduction

The human kidney is essential for life, excreting waste products by filtration of blood in glomeruli followed by the modification of ultrafiltrate by tubules. Severe kidney failure, requiring dialysis or transplantation, affects around five million people globally [1].

Congenital defects of the kidney and urinary tract are among the most prevalent types of human malformations [2], and they are the commonest cause of severe kidney failure in young children [3]. Furthermore, Wilms tumour, or nephroblastoma, is the commonest solid organ malignancy in children and it arises through abnormal kidney development [4].

The embryonic metanephros is the precursor of the mature mammalian kidney [5, 6] and in humans it arises at five weeks of gestation [7]. Its two epithelial lineages derive from intermediate mesoderm. The metanephric mesenchyme differentiates into nephron epithelia comprising glomerular podocytes and Bowman capsule, and proximal and distal tubules, while the ureteric bud undergoes serial branching to form collecting ducts, each fusing with a distal tubule. Layers of nephrons are generated between seven and 34 weeks of human gestation [7]. Native mammalian kidneys also contain vessels [8, 9]. Blood endothelia derive from a combination of angiogenic invasion from outside the metanephros combined with *in situ* vasculogenesis from metanephric mesenchyme [8]. Blood vessels populate interstitial spaces between metanephric tubules and invade tufts of glomerular podocytes to form capillary loops, essential to supply blood for filtration [8, 10, 7].

Tissue resident macrophages promote development and function of a range of organs [11]. Developing murine metanephric kidneys contain macrophages [12–15]. In animal experimental models, various roles have been attributed to macrophages including clearance of apoptotic cells [12, 13] and enhancing vessel morphogenesis [15]. Developing human kidneys also contain macrophages [16–19], but their possible roles there are unclear, in large part because of the challenges of undertaking functional studies on human fetal tissues.

Organoids derived from pluripotent stem cells (PSCs) are being used to model normal human kidney development [20–23] and early onset genetic diseases [24, 25]. Human PSC (hPSC)-derived nephron organoids are rich in glomerular podocytes and tubules, reflecting their *in vitro* derivation from intermediate mesoderm-like precursors, as occurs during native embryogenesis [26, 27]. These organoids also contain endothelia between tubules [23] but during *ex-vivo* organoid culture they rarely invade glomeruli to form capillary loops, as will occur once implanted *in vivo* [21, 28]. PSC-derived nephron organoids do not contain macrophages or express sets of genes that characterise these innate immune cells [22, 29, 25]. Indeed, the lack of macrophages is expected given that during normal development kidney macrophages originate from haematopoietic precursor cells residing within yolk sac and liver [30, 31], lineages not generated in nephron organoid culture.

The generation of macrophages from hPSCs has been described [32]. We hypothesised that combining hPSC-derived maturing macrophages with hPSC-derived nephron precursors would modify kidney development in organoid cultures. Here, we explored the possible effects of adding different numbers of macrophages harvested at earlier or later stages of their *in vitro* maturation.

## Methods

### Human Tissues

Human embryonic tissues were collected after maternal consent under ethical approval (18/NE/0290 and 23/NE/0135) and were provided by the Medical Research Council and Wellcome Trust Human Developmental Biology Resource (http://www.hdbr.org/). Fetal kidneys were fixed, processed, and paraffin sections prepared as described [33, 23].

### Human PSCs

MAN13, a human embryonic SC line that we have extensively used to generate nephron organoids [21, 23, 25], was grown on 6-well plates coated with 5 mg/mL recombinant human Vitronectin (Life Technologies, A14700) in TeSR1 (STEMCELL Technologies, 85850). Cells were passaged with 0.5 mM EDTA solution (pH 8) (Invitrogen, 15575-038) and replated in TeSR1 containing RevitaCell Supplement (100X; Thermo Fisher, A2644501) for 24 hours.

### Lentiviral reporter gene transduction

MAN13 PSCs used to generate macrophages were transduced with a reporter gene to facilitate their tracking within living organoids. Lentiviral vectors transducing the *enhanced green fluorescence protein* (*EGFP*) reporter gene driven by the strong *elongation factor 1 a (EF1α*) promoter, were generated as follows. HEK-293T cells in 15 cm dishes at 50% confluency were transfected by calcium phosphate precipitation with 10 μg pRRL.sin.cppt.EF1α-EGFP-WPRE [21], 10 μg pMDLg-pRRE, 3.4 μg pMD2.G, and 2 μg pRSV-Rev per dish. Cell media was collected over two days and centrifuged at 6,000 x g overnight, the pellet resuspended in PBS, and this suspension centrifuged in a SW40-Ti rotor (Beckman Coulter Ltd, High Wycombe, UK) at 50,000 x g for 90 minutes. The pellet was resuspended in PBS, at 1:2,000 of the original total volume of cell media that had been collected. The viral vector titre was calculated by fluorescence-activated cell sorting detecting EGFP fluorescence in HEK-293T cells transduced with serial dilutions of the vector, then determining the percentage of EGFP positive cells. MAN13 cells were transduced at a multiplicity of infection of 5 and were incorporated into nephron organoids, as described below.

### Production of macrophages from human PSCs

The method was adapted from a published protocol [32]. One well in a six-well plate of confluent MAN13 cells was fed with fresh TeSR1 media supplemented with 50 ng/ml BMP4 (R&D, 314-BP-010), 50 ng/ml VEGF (R&D, 293-VE-010) and 20 ng/ml SCF (Thermo Fisher, PHC2115). Cells were passaged using the StemPro EZPassage tool (Thermo Fisher, 23181010) and embryoid bodies (EBs) were generated in suspension plates (Grenier, 657970) for four days, supplemented with cytokines on day two. EBs were collected and transferred to six-well plates (10-15 EBs per well) coated with 0.1% w/v porcine gelatin (Sigma, G9136). EBs were fed with X-VIVO15 (Lonza, BE02-060F) media supplemented with 100 ng/ml CSF1 (BioLegend, 574806), 25 ng/ml1 IL-3 (Peprotech, 200-03-10), 2 mM Glutamax (Gibco, 35050061), 0.05 mM β-mercaptoethanol (Gibco, 31350-010), and this media was replaced every three to four days. After two to three weeks, EBs started to shed non-adherent hematopoietic stem cells (HSCs) into suspension. These cells were harvested from the supernatant and plated into six-well plates in X-VIVO15 media supplemented with 100 ng/ml CSF1 (BioLegend, 574806), 2 mM Glutamax (Gibco, catalogue number 35050061) and maintained for up to 11 days during which period they underwent macrophage differentiation.

### Differentiation of PSCs into human nephron organoids

PSCs were induced to form organoids over 25 days with a seven-day 2D phase followed by an 18-day 3D phase, as described [21, 23, 25]. MAN13 cells were plated into a well of a six-well plate coated with 5 μg/mL recombinant human Vitronectin (Life Technologies) and fed with TeSR1 (STEMCELL Technologies, 85850) media supplemented with RevitaCell Supplement (100X; Thermo Fisher, A2644501). Differentiation was initiated the following day by replacing media with STEMdiff APEL2 (StemCell Technologies, 05270) supplemented with 1% (v/v) PFHM-II Protein-Free Hybridoma Medium (Thermo Fisher Scientific, 11370882), containing 8 μM CHIR-99021 (Tocris, 4423). APEL2 with PFMH-II is hereafter referred to as ‘basal media’. On day four of the 2D protocol, cells were fed with basal media plus 200 ng/mL FGF-9 (Peprotech, 100-23) and 1 μg/mL heparin (Sigma-Aldrich, H3149-25KU). Media was replenished every day until day seven of the 2D phase when cells were dissociated into a single cell suspension by TrypLE (Gibco, 12604-021) for 1-2 minutes at 37°C followed by centrifugation at 400 x g and then they were resuspended in basal media. The suspension was aliquoted as 2×10^5^ cells in 1.5 ml Eppendorf tubes and centrifuged at 6000-7000 rpm for two minutes. Pellets, comprising nephron organoid precursor cells, were transferred onto MilliCell culture inserts (0.4 μm pore size; Millipore, PICM03050), with three organoid pellets placed on each insert and they were pulsed with basal media plus 5 μM CHIR-99021 for one hour. Next, the media was replaced with APEL2 supplemented with 200 ng/ml FGF9 and 1 μg/mL heparin (Sigma-Aldrich, H3149). From this point onwards, media was replaced every two days. On day 12 of the 3D organoid phase, basal media without additives was added until day 18, the end of the culture period.

### Incorporation of hPSC-derived macrophages into kidney organoids

Human PSC-derived macrophages were harvested at days 6, 9 and 11 of the final stage of their differentiation protocol and they were mixed with 2×10^5^ nephron precursor cells from day 7 of the 2D differentiation phase. Most of the experiments used day 6 or day 9 macrophage cells, that we also designate as ‘earlier’ or ‘later’ stage maturing macrophages. Macrophages were added at one of three cell numbers: 2,000, 10,000 or 40,000 cells, hereafter simply called ‘1%’, 5%’ or ‘20%’, in comparison with the constant number of nephron precursor cells used to generate each organoid. EGFP expression was visualised under an inverted epifluorescence microscope (Leica) in composite organoids over the next 18 days in 3D culture.

### Immunocytochemistry

PSCs, EBs and maturating macrophages were fixed with 4% PFA flowed by PBS washes. Blocking was undertaken with 1% bovine serum albumin (BSA; Sigma-Aldrich, 9048-46-8) with 10% serum from the species in which the secondary antibody was raised. Primary antibodies (Table 1) were incubated overnight at 4°C. Fluorescent secondary antibodies were incubated during 45-60 minutes at room temperature followed by 4′,6-diamidino-2-phenylindole (DAPI) nuclear staining and PBS washes. Images were acquired on an Olympus IX-71 inverted fluorescence microscope and captured using a QImaging Retiga SRV camera [25].

**Table 1.** List of primary and secondary antibodies.

| Primary Antibody | Host | Supplier | Catalogue | Application | Dilution |
| --- | --- | --- | --- | --- | --- |
| PECAM-1(CD31) | rabbit |  | Ab176533 | Immunohistochemistry<br>(fluorescent) | 1:200 |
| PECAM-1(CD31) | mouse | Cell<br>Signaling | 3528<br>(89C2) | Immunohistochemistry<br>(chromogenic) | 1:100 |
| Synaptopodin | mouse | Santa Cruz | sc-50459 | Immunohistochemistry<br>(chromogenic) | 1:200 |
| CD45 | rabbit | Thermo<br>Fisher | A700-012<br>(BL-178-12C7) | Immunohistochemistry<br>(fluorescent) | 1:250 |
| CD68 | mouse | Abcam | Ab201340 | Immunohistochemistry<br>(fluorescent) | 1:200 |
| CD74 | rabbit | Abcam | Ab270265 | Immunohistochemistry<br>(fluorescent) | 1:200 |
| CD81 | mouse | Novus<br>Biologicals | NBP1-44861 | Immunohistochemistry<br>(fluorescent) | 1:200 |
| 25f9 | mouse | Santa Cruz | SC-52698 | Immunohistochemistry<br>(fluorescent) | 1:200 |

| Secondary<br>antibody | Host | Supplier | Catalogue | Application | Dilution |
| --- | --- | --- | --- | --- | --- |
| Alexa Fluor 594 | goat -<br>anti<br>rabbit | Thermo<br>Fisher | A32740 | Immunohistochemistry<br>(fluorescent) | 1:200 |
| Alexa Fluor 594 | goat -<br>anti<br>mouse | Thermo<br>Fisher | A11032 | Immunohistochemistry<br>(fluorescent) | 1:200 |
| Biotinylated<br>antibody | Goat<br>anti-<br>rabbit | VectorLabs | BA-1000 | Immunohistochemistry<br>(chromogenic) | 1:200 |
| Biotinylated<br>antibody | Goat -<br>anti<br>mouse | VectorLabs | BA-9200 | Immunohistochemistry<br>(chromogenic) | 1:200 |

### Immunostaining of histology sections of organoids and native human fetal kidneys

Organoids were fixed in 4% paraformaldehyde, embedded in paraffin and sectioned at 5 µm. Organoid and native human kidney sections were dewaxed and rehydrated, and slides were boiled in a microwave in 10 mM sodium citrate buffer (pH 6.0; Sigma-Aldrich, C9999). After cooling to room temperature, endogenous peroxidase activity was blocked using 0.3% hydrogen peroxide (Sigma-Aldrich, H1009) in PBS for 10 minutes. Sections were permeabilized using 0.2%Triton X-100 (Sigma-Aldrich) for 10 minutes and blocked using 1% BSA (Sigma-Aldrich 05470) with 10% serum from the species in which the secondary antibody was raised. Sections were incubated overnight at 4°C with the primary antibody diluted in 1% BSA in PBS. Primary antibodies were omitted for negative controls. Biotin-conjugated species-specific secondary antibodies with 1% BSA were incubated at room temperature for 2 hours. Following PBS washes, slides were incubated in avidin-biotin enzyme complex (Vector Laboratories, VECTASTAIN Elite ABC Reagent, PK-6100) for one hour at room temperature. Peroxidase activity was detected with SIGMAFAST 3, 3 diaminobenzidine peroxidase substrate solution (Sigma-Aldrich, D4293) and sections were counterstained with haematoxylin (Sigma-Aldrich, HH516). Sections were dehydrated and mounted with DPX mounting medium (Fisher Chemical, D/5330/05). Brightfield images were acquired on a 3D-Histech Pannoramic-250 microscope slide-scanner using a x20 objective (Zeiss) and selected images were captured using the Case Viewer software (3D-Histech).

For immunofluorescence on paraffin sections, tissues were dewaxed and rehydrated as described above. Blocking was undertaken with 1% BSA (Sigma-Aldrich, 9048-46-8) with 10% serum from the species in which the secondary antibody was raised. Primary and Alexa Fluor secondary antibodies are listed in Table 1. To block autofluorescence, Vector® TrueVIEW® kit (Vector, SP-8500) was used following user guide recommendations. Nuclei were stained with DAPI and mounted with mounting medium from the same kit.

### Quantification of cells and tissues within organoids

For macrophage quantification, sections of kidney organoids were immunostained for CD68 [16]. CD68 positive cells were counted by randomly selecting four x40 objective histology fields of transverse sections (two from the glomerulus-rich “cortex” and two from the tubule-rich “medulla”) of each organoid. The quantity for each organoid was defined as the average number of positive cells in the four sections. To quantify endothelia, histology sections through the maximum transverse diameter of the organoid were immunostained for CD31/platelet endothelial cell adhesion molecule-1 (PECAM-1). The positive stained area was measured using Image J and expressed as a proportion of the total area. For quantification of the ‘mass’ of glomerular epithelia, similar analyses were undertaken on histology sections immunostained for synaptopodin. For quantification of glomeruli containing endothelia, twenty randomly selected glomeruli per organoid were imaged. A glomerulus was designated as positive for vessels if it contained at least three CD31/PECAM-1 positive isolated cells, or an overt blood vessel, or both features. Statistical analyses were undertaken using GraphPad Prism 10.2.2. Results were expressed as mean±SEMs for parametric data, or medians and interquartile ranges for non-parametric data sets. Parametric data sets were analysed using one-way ANOVA for multiple comparisons followed by *post-hoc* t tests. Non-parametric data were analysed by Kruskal Wallis tests followed by Mann-Whitney U-tests.

## Results

### Differentiating MAN13 hPSCs to macrophages

hPSCs were transduced with lentiviral vector carrying a GFP expression construct. To differentiate them to macrophages, cells were fed with a series of media containing specific growth factors (Figure 1a), using a published protocol [32]. hPSCs were differentiated through an EB stage during which haematopoietic stem cells (HSCs) emerged in suspension. These shed cells were harvested and subsequently matured as adherent cells over a further 9 to 11 days *in vitro* to generate macrophages. Figure 1b shows phase contrast images of this sequence, and Figure 1c depicts the sequence visualised under fluorescence showing extensive GPF expression. Cells were investigated using immunocytochemistry, with antibodies against HSC and macrophage markers (Figure 2) [16, 17, 34, 32]. PSCs were negative for macrophage markers. At the start of the final 11 days of the maturation protocol, cells were positive for the HSC marker CD45 and they showed increasing positivity for the mature macrophage marker 25f9 over the next 11 days. Maturing macrophages were also positive for CD74 and CD81, markers of resident kidney macrophages [34], and for CD68, as found in human fetal kidneys [16, 17].

**Figure 1.**
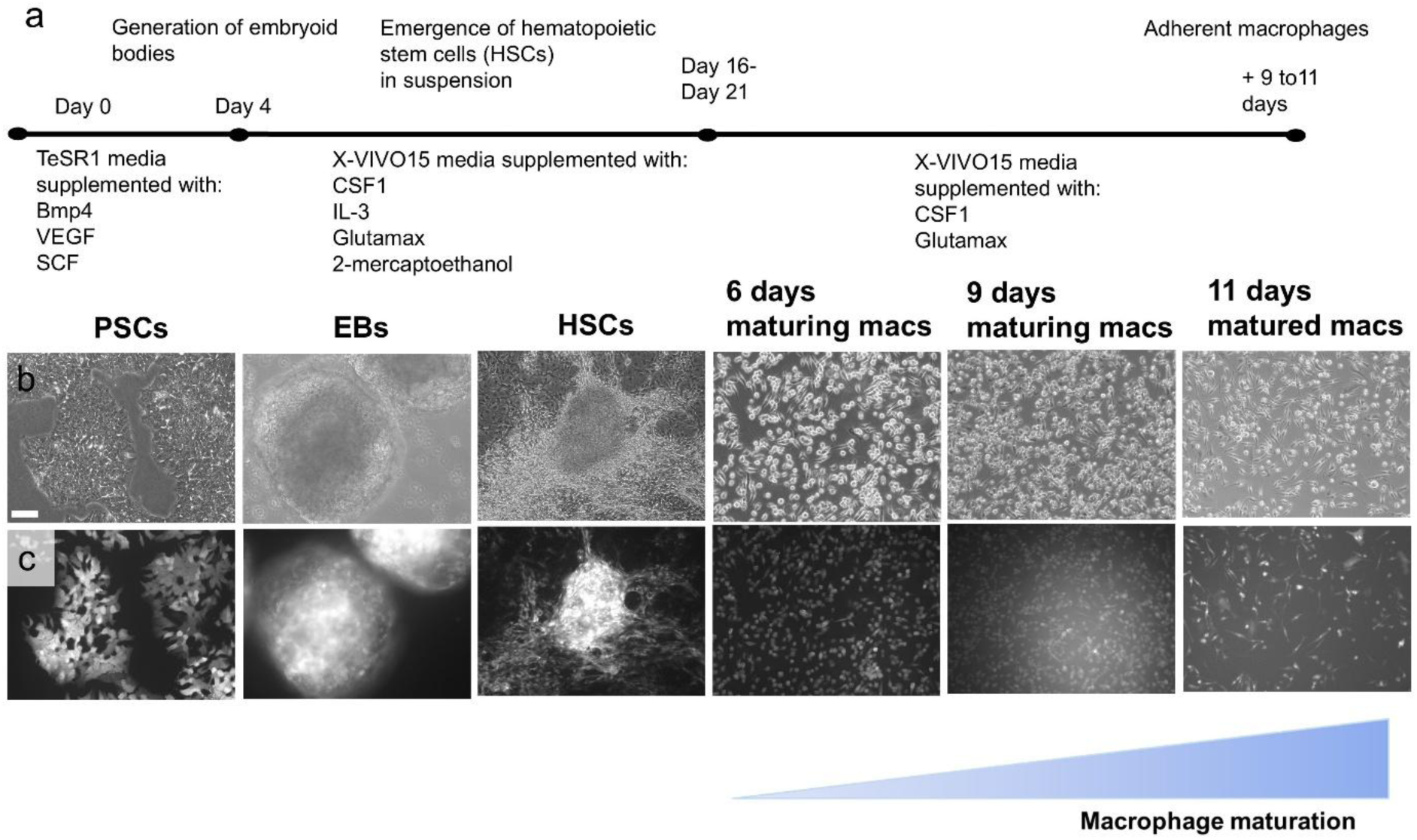
Deriving macrophages from hPSCs. **a.** Graphic of the protocol showing sequential exposure of cells to specific media and growth factor supplements. SCs form EBs over the first four days. HSCs are shed over the next 12 to 17 days. Adherent macrophage precursors mature over the final 11 days of the protocol. **b.** Phase contrast images showing hPSCs differentiated sequentially into EBs, shed HSCs and finally maturing macrophages. **c.** The same sequence as b., showing widespread expression (white) of the EGFP reporter gene as detected by fluorescence microscopy. Bar is 75 µm.

**Figure 2.**
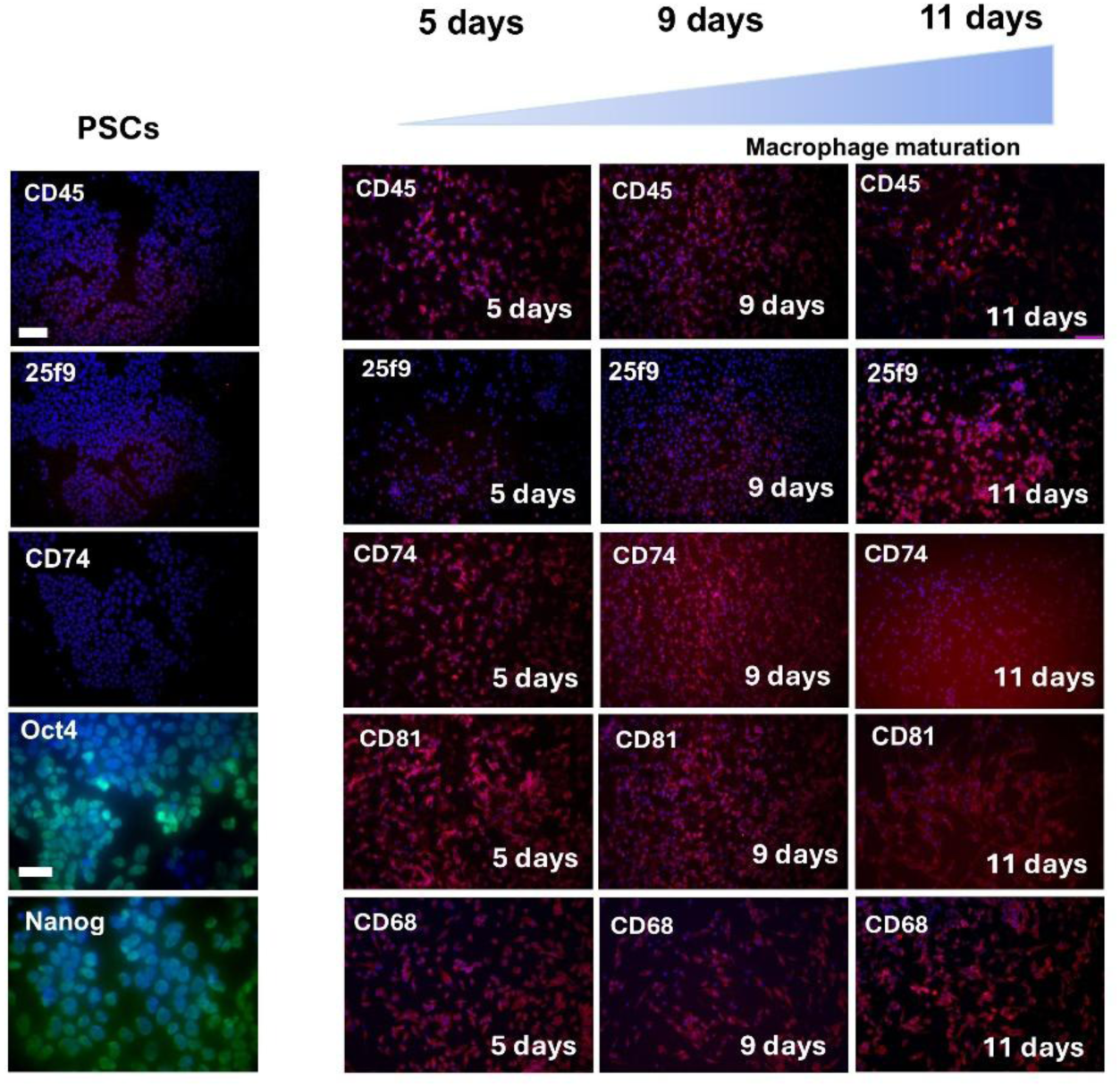
Fluorescence immunocytochemistry of hPSCs and maturing macrophages. Cell nuclei were stained (blue) with DAPI. Cells were immunostained for HSC and macrophage markers (red) or pluripotency markers (green). Bar is 100 μm apart from 40 μm for pluripotency marker frames. **Column 1.** PSCs were negative for HSC/macrophage lineage markers (CD45, 29f9 and CD74) but, as expected, were positive for classical SC transcription factor markers (green nuclei) Oct4 and Nanog. **Columns 2, 3 and 4.** At the start of the final 11 days of differentiation protocol, cells were positive for the HSC marker CD45 (red). Over the next 11 days, they acquired positivity for the mature macrophage marker 25f9 (red). Maturing macrophages were positive for CD74, CD81, and CD68 (all red).

### Detection of macrophages within nephron organoids

MAN13 PSCs were differentiated into organoids over a 25-day protocol (Figure 3a). Nephron precursor cells were generated over seven days in 2D culture (Figure 3b) and they were dissociated and pelleted onto tissue culture inserts. Over the next 18 days each pellet was differentiated from an amorphous cell aggregate to a structure with internal patterning. In organoids where nephron precursor cells were mixed with macrophages, these appearances were grossly preserved, as assessed by phase contrast microscopy (Figure 3c-e). Using fluorescence microscopy of these composite organoids (Figure 3f-h), positive signals for the EGFP reporter were detected over the whole of the 18-day 3D culture, suggesting that viable macrophages were present throughout this period. As nephrogenesis proceeds within organoids, the tissue becomes thicker so that visualisation of macrophages within cultures may be technically limited. Therefore, we identified macrophages using CD68 immunostaining within tissue sections of organoids harvested at the end of the 3D culture period (Figure 4). In sections of human fetal kidneys (Figure 4a), macrophages were detected in the stromal/interstitial compartment between tubules and glomeruli. Nephron organoids also contained tubules and glomeruli (Figure 4b and c). In those generated without added hPSC-derived macrophages, no CD68 positive cells were detected (Figure 4b), whereas macrophages were detected in between tubules and around glomeruli within composite organoids (Figure 4c). Next, we quantified macrophages within organoids at the end of the culture period, comparing numbers when 1%, 5% or 20% of macrophages were incorporated from earlier (day six) or later (day nine) in the final phase of the *in vitro* differentiation protocol. There was a positive correlation of numbers between macrophages counted within organoid sections and numbers added (Figure 4d and e).

**Figure 3.**
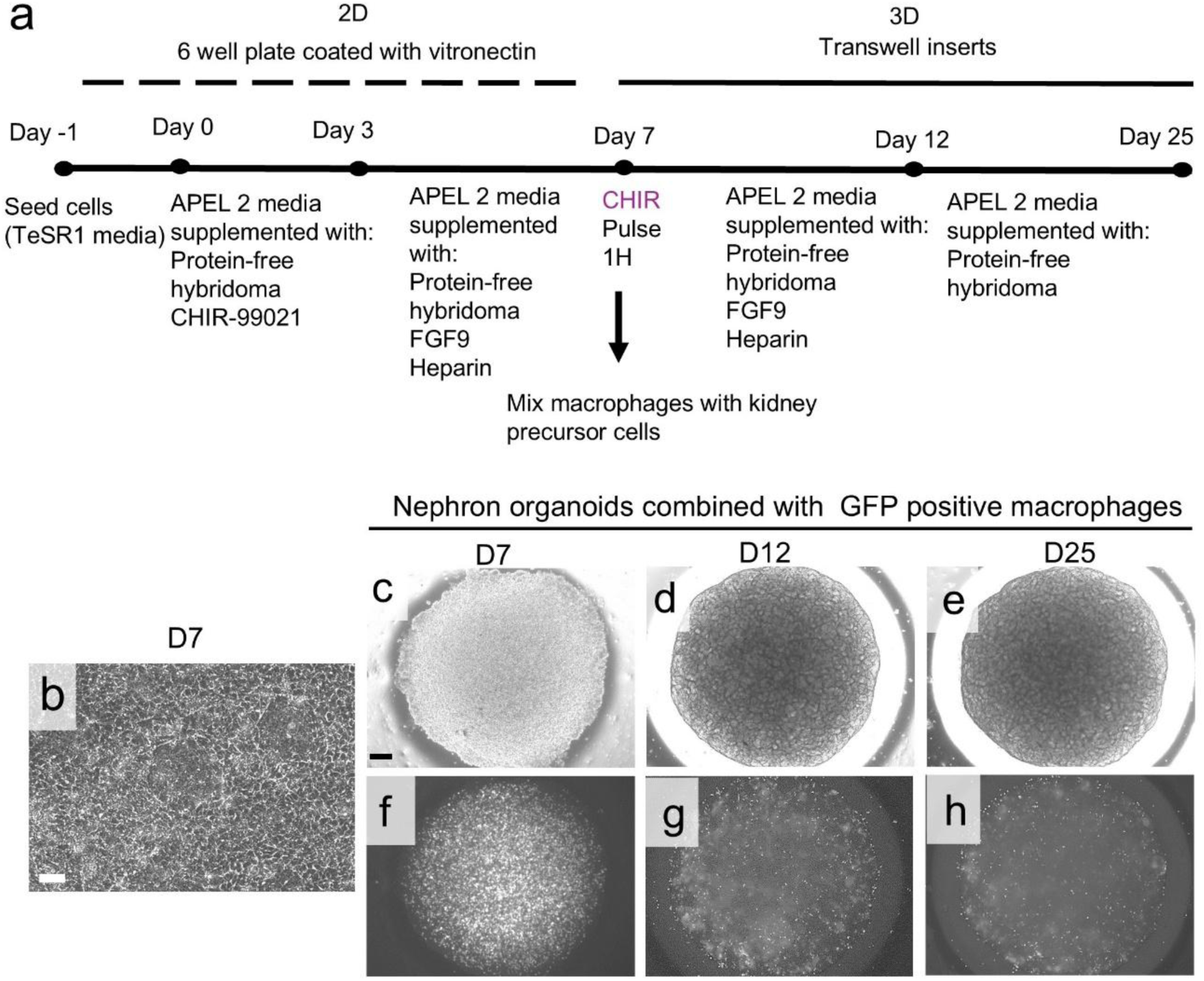
Generation of nephron organoids. **a.** The 25-day protocol whereby hPSCs were differentiated into nephron organoids. At the end of the 2D phase (*Day 7*), nephron precursor cells were dissociated and pelleted onto tissue culture inserts and maintained for a further 18 days. In some experiments, hPSC-derived macrophages were mixed with the nephron precursor pellet at Day 7. During this period, organoids are disc-shaped, with a final maximum diameter of around 0.5 cm. **b-h.** Images of cell cultures. In the sequence depicted in c-h, matured macrophages were mixed with nephron precursor cells (here, day 11 macrophages, 5% were used). **b-e.** Phase contrast images of Day 7 nephron precursor cells before they were dissociated (b). On the day when cells were placed on inserts the pellet appeared, as viewed from above, as a featureless disc-shaped aggregate (c). Over the next 19 days the cell mass acquired internal patterning (d and e). **f-h.** Fluorescent images of a composite (nephron precursor plus EGFP-transduced macrophage). Reporter gene signals were detected as discrete dots (white) throughout the 3D culture period, suggesting the presence and persistence of viable macrophages. Bars are 500 µm.

**Figure 4.**
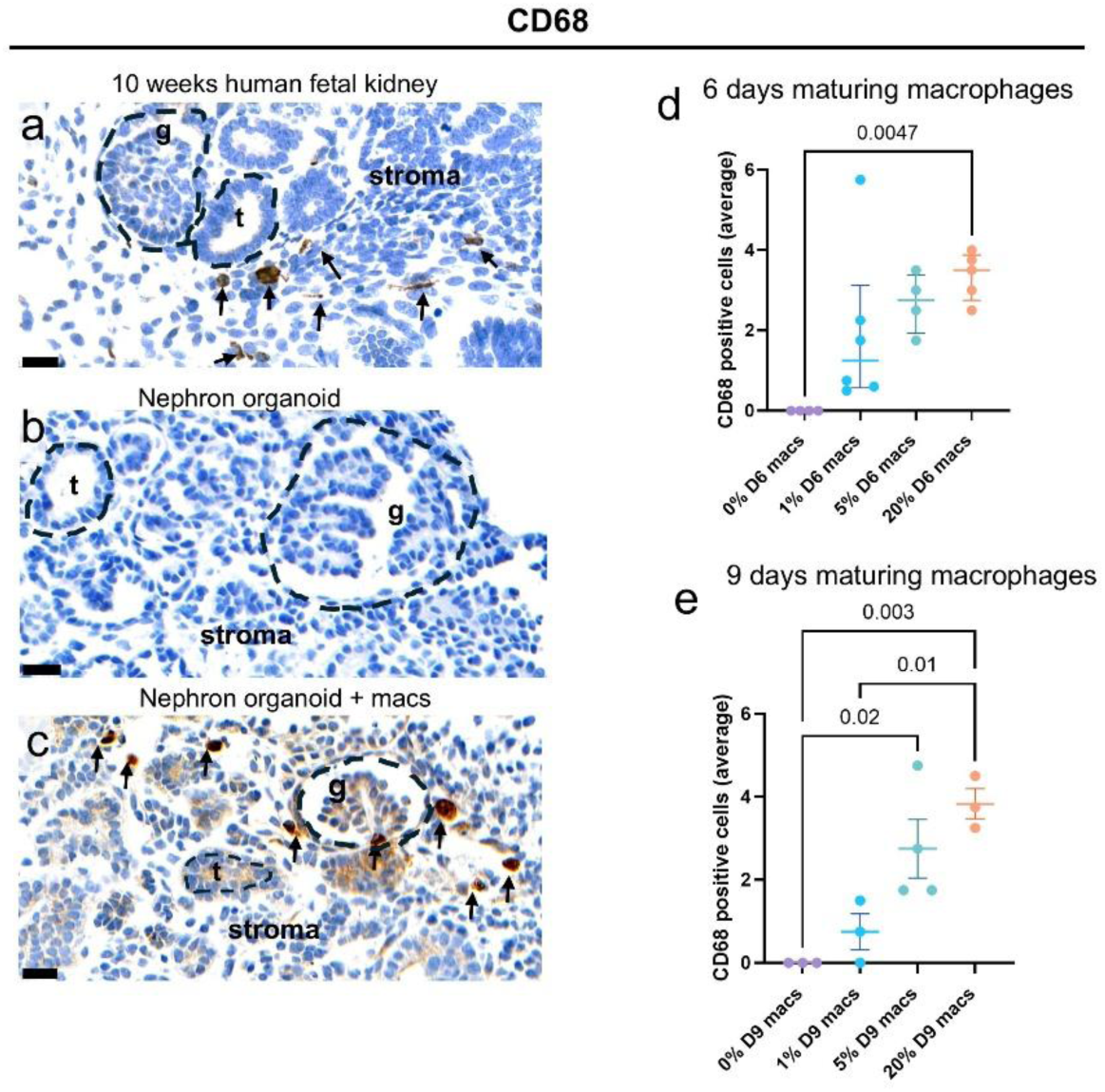
Detection of macrophages in tissue sections. **a-c.** Immunohistochemistry for macrophages marker CD68 (brown) with cell nuclei stained (blue) with haematoxylin. Bars are 20 µm. Note the CD68 positive cells (arrows) in the stromal compartment between glomeruli (g) and tubules (t) in a native human fetal kidney at 10 weeks of gestation (**a**) and in the composite (nephron precursor + macrophage) organoid (**c**). By contrast, the organoid made from only nephron precursor cells (**b**) did not contain CD68 positive cells. **d and e.** Quantification of CD68 positive cells per 0.12 mm^2^ at the end of the 3D culture period after adding either early (6 days; d) or later (9 days; e) maturing macrophages. Note positive correlations between macrophage numbers added at the start of the 3D culture and counts within histology sections of organoids harvested at the end of the 18 day 3D culture. Each dot is one organoid. Data in d are shown as median and interquartile range, while data in e are shown as mean±SEM. P values are specified when <0.05

### Analyses of vessels and glomeruli within organoids

Histology sections of composite organoids were co-immunostained for CD31/PECAM-1 and CD68. Networks of vessels were detected, and macrophages were noted to sometimes be in proximity to vessels positive for CD31/PECAM-1 (Figure 5a). In brightfield imaging, CD31/PECAM-1 positive cells, often taking the form of a vessel with an apparent lumen, were prominent in interstitial spaces of nephron organoids generated either without (Figure 5b) or with (Figure 5c) added macrophages. We then quantified vessels in organoids using two approaches. First, the proportional area occupied by vessels in sections of organoids was measured. In organoids without added macrophages, around 1% of the total area was occupied by vessels and this value did not change significantly in organoids generated with 1%, 5% or 20% day 6 (early stage) macrophages (Figure 5d). In similar experiments where day 9 (later stage) macrophages were used, there appeared to be a concentration-dependent effect, with 1% added macrophages reducing the total vessel area compared with 5% added macrophages (Figure 5e). Second, we assessed whether glomeruli contained endothelia. In all conditions studied, most glomeruli imaged were negative (Figure 5f) but rare glomerular tufts contained CD31/PECAM-1 positive cells manifesting as capillary-like structures or as isolated positive cells (Figure 5g). Quantification showed no significant change in proportions of positive glomeruli in organoids with added macrophages (Figure 5h and i). Although a trend to increased numbers was noted when early-stage macrophages were added, this did not attain statistical significance. Next, we assessed the ‘mass’ of glomeruli by quantifying the percentage area of tissue sections occupied by cells immunostained for synaptopodin. Podocyte tufts were positive in both organoids generated without (Figure 6a) or with (Figure 6b) added macrophages. In experiments where early stage (day 6) macrophages were added, there was a statistically significant increase in glomerular area, by approximately three-fold, when nephrons were generated with 5% or 20% added macrophages (Figure 6c). By contrast, when organoids were generated with more mature (day 9) macrophages, 1% added macrophages were found to have reduced the glomerular area compared with 5% added macrophages (Figure 6d).

**Figure 5.**
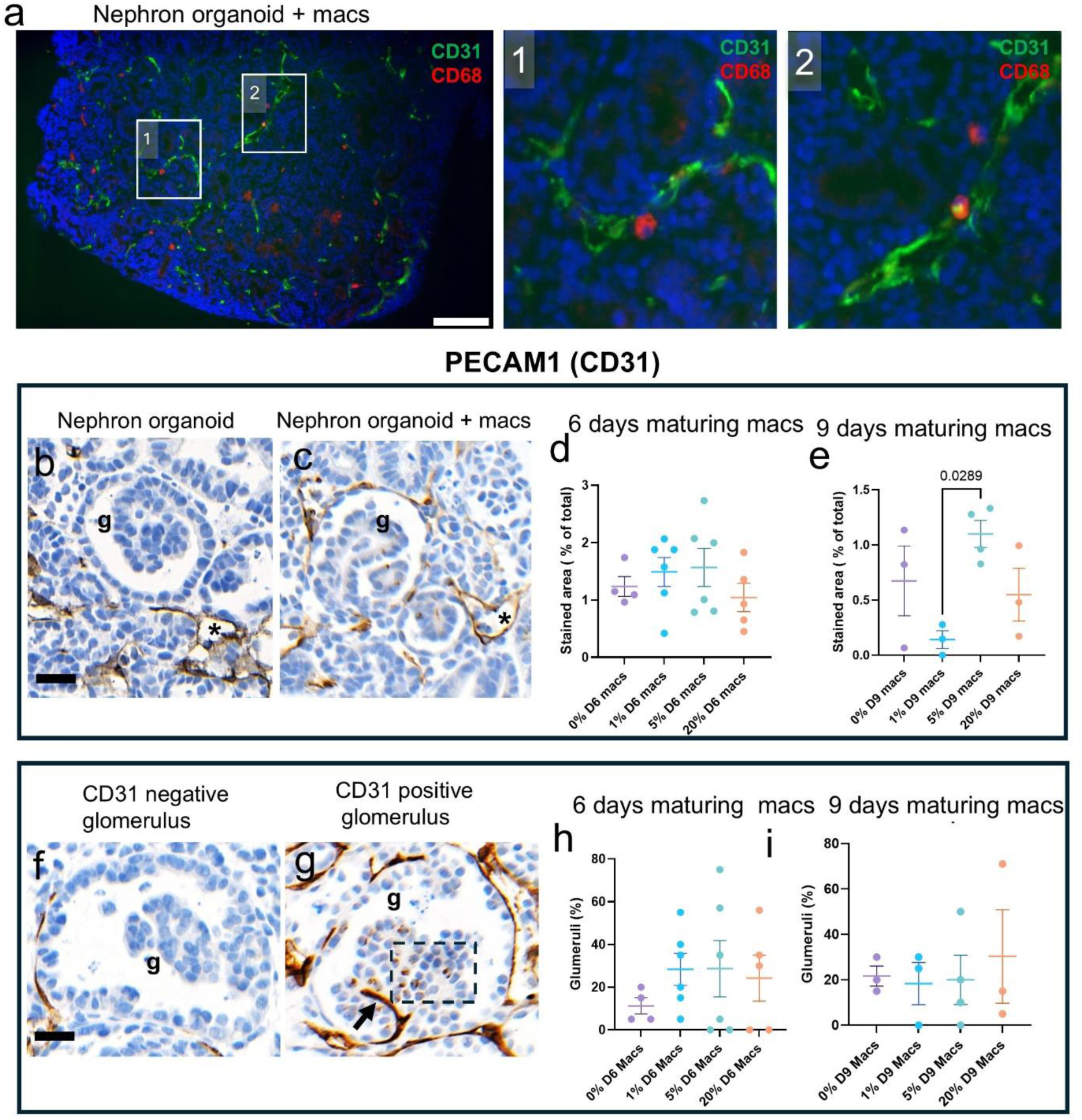
Vessels within nephron organoids. **a.** Fluorescence image of a composite organoid. Dual immunostaining of blood vessels (CD31/PECAM-1; green) and macrophages (CD68; red) revealed macrophages, sometimes in proximity vessel networks. Nuclei counterstained with DAPI (blue) and bar is 100 µm. Middle and right frames are enlarged views of boxes 1 and 2. **b and c.** Brightfield images of organoid histology, with nuclei stained with haematoxylin (blue), showing CD31/PECAM-1 positive cells (brown), sometimes forming patent (*asterisks*) vessel-like structures, in interstitial spaces of nephron organoids generated without (b) or with (c) macrophages. **d.** Areas occupied by vessels was unaltered in composite organoids made with (*day 6*) early stage macrophages. **e.** Addition of later stage (*day 9*) macrophages showed that 1% added cells reduced total vessel area compared with 5% macrophages**. f and g.** Brightfield images of glomeruli in organoids, with nuclei stained with haematoxylin (blue), showing an example lacking endothelia (f) compared with a glomerulus (g) containing CD31/PECAM-1 positive cells (brown) appearing as a vessel-like structure (*arrow*) or isolated cells (*box*). Bars are 20 µm. **h and i.** Added macrophages did not significantly alter the percentage of positive glomeruli, although a positive trend was noted with day 6 macrophages.

**Figure 6.**
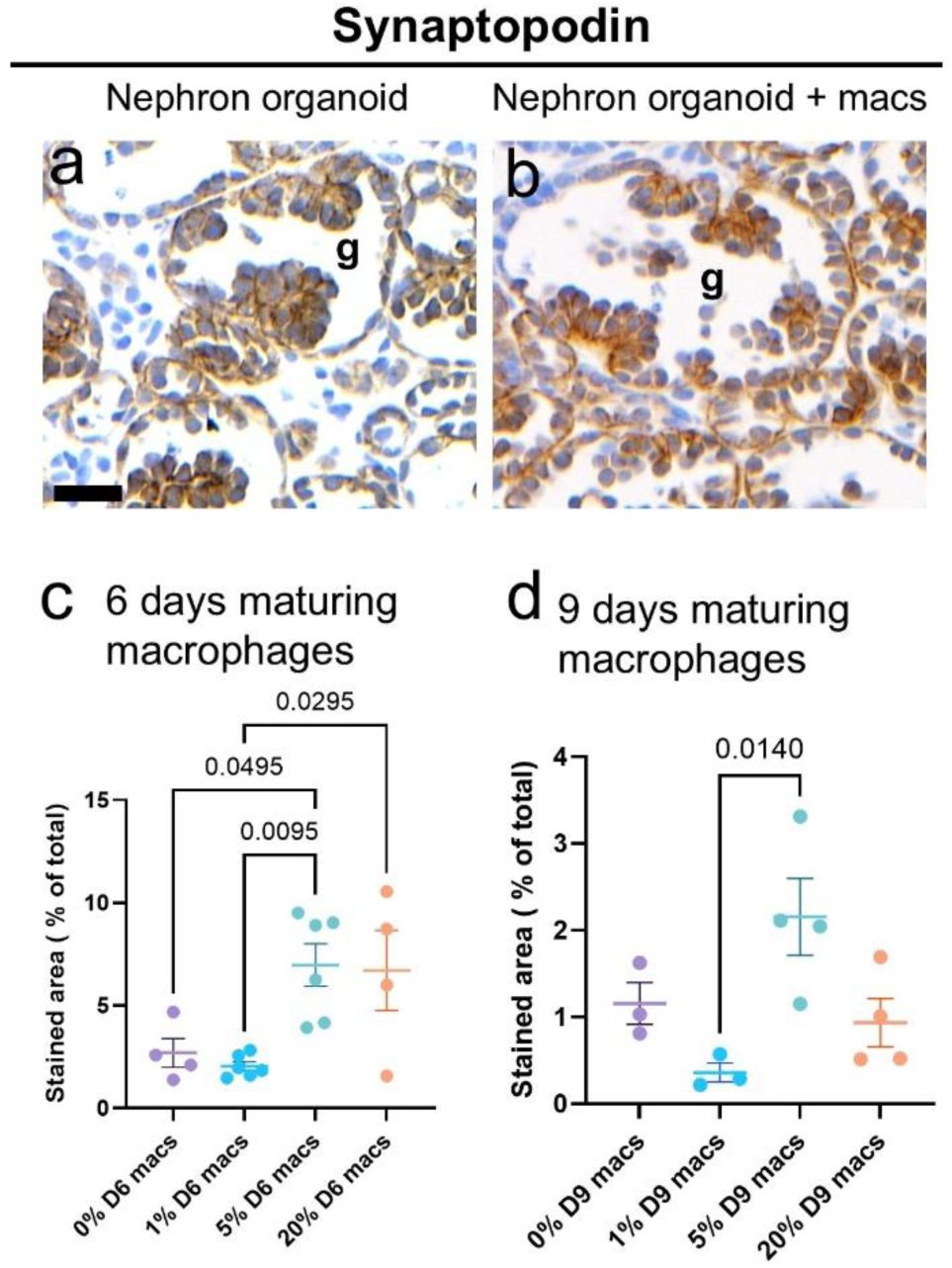
Synaptopodin immunostaining a and. **b.** Brightfield histology images, with nuclei stained with haematoxylin (blue), showing synaptopodin positive cells (brown) in glomerular tufts of organoids generated without (a) or with (b) added macrophages. Bar is 20 µm. **c and d.** Percentage areas of histology sections positive for synaptopodin immunostaing. Addition of 5% day 6 maturing macrophages significantly increased glomerular area compared with organoids without macrophages (c). In organoids generated with day 9 macrophages there was a significant increase comparing 1% and 5% macrophages. Data are mean±SEM, analysed by one-way ANOVA with multiple comparisons followed by post hoc tests. Each dot represents one organoid.

### High numbers of macrophages inhibit organoid growth

Transverse paraffin sections taken at the mid-height of each organoid were imaged (Figure 7a and b) and their areas quantified (Figure 7c and d). In experiments where 1% or 5% day 6 or day 9 maturing macrophages were added to nephron precursors, the final organoid areas were not significantly different to organoids generated without macrophages. In contrast, using 20% of either earlier or later maturing macrophages, organ sizes were significantly less than in organoids generated without macrophages. Moreover, addition of 20% day 9 maturing macrophages was associated with disruption of tissue morphology within the medulla of organoids manifesting as prominent stromal-like and immature blastema-like cells (Figure 7e-j). Such atypical tissue was not seen in organoids generated with lower numbers of either early or late stage maturing macrophages.

**Figure 7.**
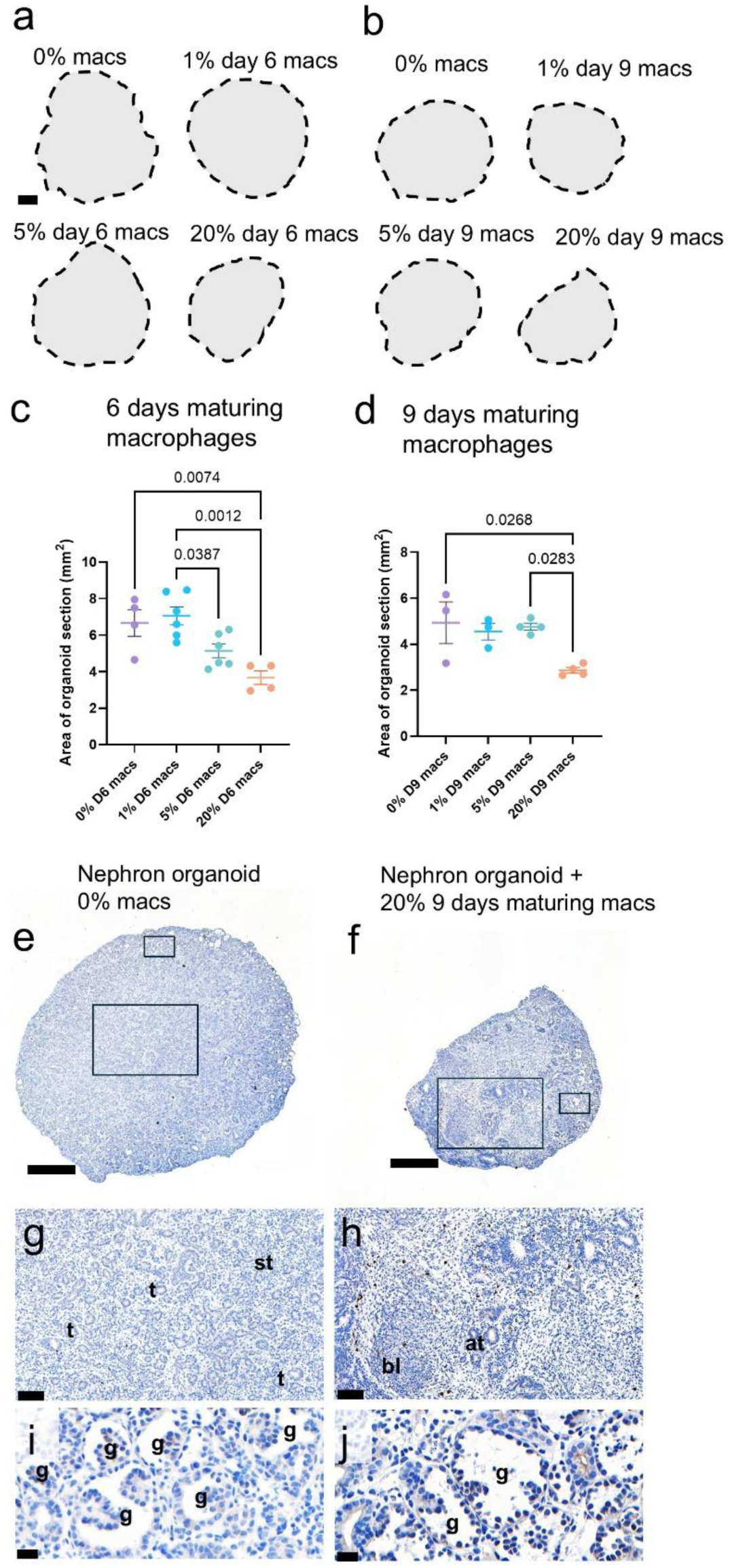
High numbers of macrophages disturb organoid growth. a and b. Outlines of histology transverse sections through the mid-height of each organoid. Scale bar is 500 μm **c and d**. Quantification of areas. Areas of organoids made with 20% added earlier (day 6) or later (day 9) maturing macrophages, were significantly less than organoids generated without macrophages. **e-j.** Histology of organoids with nuclei stained blue and CD68 macrophages immunostained brown. An organoid generated without added macrophages is shown in low power (e), with enlarged boxes showing that its medulla (g) is dominated by tubules (*t*) and its cortex (i) dominated by glomeruli (*g*). Low power of a histology section of an organoid generated with 20% day 9 maturing macrophages (f). Its medulla (h) contained stomal-like (s) and blastema-like (b) tissues with nearby macrophages (brown), while glomeruli are present in its cortex (j). Bars are 500 μm in e and f, and 20 μm in g-j.

## Discussion

Macrophages have been detected in the healthy developing human metanephric kidney [13, 19] but their possible roles are unknown. To test the hypothesis that macrophages modulate human kidney development, we harnessed hPSC technology, combining hPSC-derived macrophages with hPSC-derived kidney precursors. The current experiments demonstrate that it is technically feasible to populate hPSC derived-nephron organoids with hPSC-derived maturing macrophages. This platform provided the opportunity to explore the possible effects of macrophages on early human kidney development in a controlled *in vitro* setting. At the end of the 18-day 3D culture period, macrophages, assessed by CD68 immunostaining, were not detected in organoids generated without added macrophages, supporting RNA sequencing analyses of hPSC-derived kidney organoids from several laboratories [22, 29, 35, 25]. In contrast, organoids in which hPSC-derived macrophages had been added contained CD68 positive cells in interstitial spaces, as did human fetal kidneys. These observations are consistent with the fact that nephron organoids are generated *via* intermediate mesoderm-like precursor cells, as occurs *in vivo* [27], whereas *in vivo* macrophages populate the metanephros after being generated by other, distant haematopoietic cells [30, 31]. Quantification of macrophages within histology sections at the end of the organoid culture period showed a positive correlation with the number of macrophages added. This, together with the detection of EGFP-expressing cells within living composite organoids, confirmed that added macrophages survive in the organoid environment and thus the robustness of our composite nephron/macrophage organoid model.

The proximity of macrophages to vessels in organoids from the current experiments resembled the pattern reported in the mouse metanephric kidney studies [15]. This, together with murine observations that macrophages enhance vessel patterning in the murine metanephric explants [15], prompted us to explore possible vascular effects of adding macrophages in composite human nephron organoids. Added macrophages, however, neither increased overall CD31/PECAM-1 positive vessel area nor endothelial invasion into glomeruli, which generally remained avascular. While our results do not exclude more nuanced effects on vessels within organoids, such as modifying vascular anastomoses [15] or specific patterns of vessels, our observation show that macrophages are not advantageous for the generation and/or maintenance of vessels within interstitial spaces of human nephron organoids. *In vivo*, the formation of capillary loops within glomeruli is driven by an angiogenic growth factor gradient originating in glomerular endothelia [10]. In nephron organoid culture, this gradient appears absent or insufficient [20, 36, 21], although it can be restored in human glomeruli that continue to differentiate after subcutaneous implantation of hPSC-derived kidney progenitors into SCID-Beige mice [21]. After implantation, developing kidney organoids become perfused by the host’s blood supply and the human glomeruli acquire capillaries [21], also observed after implantation of organoids under the kidney capsule [28]. In other contexts, macrophages have been implicated in enhancing lymphatic differentiation, either by virtue of secreting growth factors or through macrophage/lymphatic transdifferentation [37], with both cell types expressing the hyaluronan receptor 1, LYVE-1 [9]. *In vivo*, fetal kidneys contain lymphatic vessels [9], and it remains possible that resident macrophages are required for their differentiation.

A previous study reported that addition of CSF-1, a macrophage trophic factor, enhanced the growth of rodent metanephric explants, increasing nephron numbers [14]. While the mechanism was unclear, it was postulated to be linked to an observed upregulation of the M2 macrophage subtype response gene *triggering receptor expressed on myeloid cells 2* [14]. Conversely, in another study of explanted rodent fetal kidneys, addition of tumour necrosis factor α (TNFα), an inflammatory cytokine, increased macrophage numbers and impaired nephron formation [13]. In the current human study, we aimed to generate resident macrophages and did not determine if they were activated to either an inflammatory M1, or the alternatively activated M2, phenotype [38].

We found that the addition of early-stage maturing macrophages significantly increased the proportional area occupied by glomeruli, as assessed by immunostaining for synaptopodin, a marker of podocytes. While the mechanism underlying this effect requires further study, we postulate that macrophages release growth factors that either directly or indirectly enhance podocyte growth. Interestingly this effect was found only with macrophages differentiated to day 6, while later stage macrophages did not have this effect.

We found that the size of organoids, as measured by the area, was not changed by the addition of low (1%) or medium (5%) numbers of added macrophages. In contrast, the addition of high number (20%) of macrophages, either from earlier or later in their *in vitro* differentiation from hPSCs, significantly reduced the area of organoids compared with those without added macrophages. Importantly, in all conditions, the starting number of nephron precursor cells was the same. While the numbers of macrophages used here cannot easily be equated with numbers resident in native metanephric kidneys, likely to be less than a few percent of all cells, the results suggest that macrophages may have harmful effects on growth when present in high numbers. This was emphasised by the additional observation that high number day 9 maturing macrophages were also associated with disrupted tissues in the middle of the organoid, with prominent stromal-like and immature blastema-like cells that were not apparent in other conditions. These aberrant cells at least superficially resemble the immature and disorganised tissues found in certain diseases of human kidney development. This includes causing dysplastic kidney malformations [16], where kidney contain incompletely differentiated tubules. Moreover, malformed human kidneys contain foci of macrophages and T lymphocytes, and TNFα is detected in the urine of human fetuses with malformed kidneys [16]. It is now appreciated that some cases of dysplastic kidneys are caused by mutations of genes involved in nephrogenesis, and aspects of the *in vivo* phenotype can be modelled in nephron organoids generated from mutant hPSCs [25]. In future, it may be informative to assess whether addition of hPSC-derived macrophages modifies the abnormal phenotype in such mutant organoids. Macrophages are also prominent in tissue sections of Wilms tumour, or nephroblastoma [39–41]. Here, M2 macrophages have been associated with a poor prognosis, partly through enhancing epithelial to mesenchymal transition of tumour cells, a prelude to metastasis [41].

The current experiments can be compared with other studies that considered potential interactions of inflammatory cells and kidney organoids. One such reported the indirect co-culture of human kidney organoids with human peripheral blood monocytes or PSC-derived macrophages [42]. Native monocytes or the PSC-derived macrophages each decreased CHIR99021-induced apoptosis in kidney precursors at the initiation of the 3D organoid protocol [42]. Moreover, native monocyte cocultures promoted expression of transcripts associated with nephron differentiation, including that encoding nephrin, a glomerular marker [42]. These observations complement our current work in which hPSC-derived macrophages incorporated into nephron organoids enhanced the percentage glomerular area. In another study [43], co-culture of human peripheral blood mononuclear cells with human PSC-derived kidney organoids induced fibrosis within the composite tissues. In this experiment, however, a variety of native inflammatory cell types were employed, rather than hPSC-derived early differentiating macrophages as described in the current study.

In summary, the current experiments show that, depending on their maturation stage and quantity, these macrophages can have beneficial or harmful effects on the growth and differentiation of human nephron organoids. Collectively, these observations support the proposition that, *in vivo*, innate immune cells play diverse roles in normal and abnormal development of native human kidneys. The mechanisms whereby macrophages have these effects requires further study.

## Acknowledgements

We thank Prof James Uney, University of Bristol, for provision of lentiviral component reagents. We also thank Ms Nicola Bates and Dr Brenda Aguero Burgos, University of Manchester, for technical contributions.

## Author Contributions

SJK, FML and ASW conceived and designed the experiments, and drafted the manuscript. FML undertook the bulk of the experiments. IB and AS provided major technical contributions regarding lentiviral and histology experiments. All authors approved the final manuscript.

## Ethics statement

Human embryonic tissues were collected after maternal consent under ethical approval (18/NE/0290 and 23/NE/0135) and were provided by the Medical Research Council and Wellcome Trust Human Developmental Biology Resource (http://www.hdbr.org/).

## Data Availability

The full scope of the paper is included in the results section of the paper.

